# Capturing auxin response factors syntax using DNA binding models

**DOI:** 10.1101/359182

**Authors:** Arnaud Stigliani, Raquel Martin-Arevalillo, Jérémy Lucas, Adrien Bessy, Thomas Vinos-Poyo, Victoria Mironova, Teva Vernoux, Renaud Dumas, François Parcy

## Abstract

Auxin is a key hormone performing a wealth of functions throughout the plant life cycle. It acts largely by regulating genes at the transcriptional level through a family of transcription factors (TF) called auxin response factors (ARF). Even if all ARF monomers analysed so far bind a similar DNA sequence, there is evidence that ARFs differ in their target genomic regions and regulated genes. Here we use position weight matrices (PWM) to model ARF DNA binding specificity based on published DNA affinity purification sequencing (DAP-seq) data. We find that the genome binding of two ARFs (ARF2 and ARF5/Monopteros/MP) differ largely because these two factors have different preferred ARF binding site (ARFbs) arrangements (orientation and spacing). We illustrate why PWMs are more versatile to reliably identify ARFbs than the widely used consensus sequences and demonstrate their power with biochemical experiments on the regulatory regions of the *IAA19* model gene. Finally, we combined gene regulation by auxin with ARF-bound regions and identified specific ARFbs configurations that are over-represented in auxin up-regulated genes, thus deciphering the ARFbs syntax functional for regulation. This provides a general method to exploit the potential of genome-wide DNA binding assays and decode gene regulation.

## INTRODUCTION

Auxin is a key hormone in plants affecting multiple developmental processes throughout the lifecycle of the plant. Most long-term developmental auxin responses (such as embryo polarity establishment, tropisms, phyllotaxis or secondary root emergence) involve modifications of gene expression by the nuclear auxin pathway (Lavy and Estelle, 2016; Weijers and Wagner, 2016). This pathway includes a family of transcription factors (TFs) called Auxin Response Factors (ARF) (Weijers and Wagner, 2016; Leyser, 2018). In the absence of auxin, the Aux/IAA repressors bind ARF TFs and form inactive multimers thereby preventing their activity (Han et al., 2014; Korasick et al., 2015). The presence of auxin leads to the degradation of Aux/IAA proteins and therefore allows ARFs to activate transcription.

ARF proteins exist in 3 classes (A, B and C) with class A corresponding to ARF activators and B and C to ARF repressors (Finet et al., 2013). Understanding ARF biochemical properties (DNA binding specificity, capacity to activate or repress transcription, capacity to interact with partners) is important to decipher how different tissues could respond differently to the same auxin signal (Leyser, 2018). ARFs are modular proteins with several functional domains: most ARFs (except ARF3/ETTIN, ARF17 and ARF23) have a PB1 domain (previously called domain III/IV) responsible for interaction with the Aux/IAA repressors, TFs from other families and possible homo-oligomerization through electrostatic head-to-tail assembly (Nanao et al., 2014; Korasick et al., 2014; Parcy et al., 2016; Weijers and Wagner, 2016; Mironova et al., 2017). ARFs also possess a DNA binding domain (DBD) from the plant specific B3 family. The structure of this DBD has been solved for ARF5 (also called Monopteros/MP) and ARF1 revealing a B3 domain embedded within a flanking domain (FD) and a dimerization domain (DD) (Boer et al., 2014). The DD allows ARF proteins to interact as a face-to-face dimer with a DNA element called an everted repeat (ER) made of two ARF binding sites (ARFbs). ARFbs have been originally defined as TGTCTC (Ulmasov et al., 1995; Guilfoyle et al., 1998) and this knowledge was used to construct a widely used auxin transcriptional reporter, DR5 (Ulmasov et al., 1997). More recently, Protein Binding Microarray (PBM) experiments suggested that TGTCGG are preferred ARFbs and a new version of DR5, DR5v2, was built based on this cis-element (Boer et al., 2014; Franco-Zorrilla et al., 2014; Liao et al., 2015). ARFs are able to bind different ARFbs configurations in addition to ER such as direct repeats (DR) or, as recently suggested, inverted repeats (IR) (O’Malley et al., 2016). Whether ARF oligomerization through the PB1 domain contributes to binding of some ARFbs configurations such as IR or DR that are not compatible with DD dimerization has been proposed but not yet demonstrated (O’Malley et al., 2016; Parcy et al., 2016). Based on affinity measurements of interaction between ARF DBD (for ARF1 and MP) and a few ER cis-elements, it was proposed that ARFs differ by the type of ER configuration they prefer: the ARF1 repressor has a much narrower range of preferences than the MP activator (this was called the molecular caliper model) (Boer et al., 2014). However, this model was established using isolated ARF DBD lacking the PB1 domain and did not include interaction with DR and IR ARFbs configurations.

Despite the central importance of ARF TFs, models reliably predicting their DNA binding specificity are still scarce (Keilwagen et al., 2011; Mironova et al., 2014) and simple consensus sequences are often used (Berendzen et al., 2012; O’Malley et al., 2016; Zemlyanskaya et al., 2016) that hardly capture possible sequence variation within the cis-element. Recently, DNA affinity purification sequencing (DAP-seq) data have offered a genome-wide view for two full-length ARF proteins of Arabidopsis (the repressor ARF2 and the activator MP) (O’Malley et al., 2016). The DAP-seq assay is technically similar to ChIP-seq but with chromatin-free isolated genomic DNA and with a single recombinant protein added. Based on TGTC consensus sequence as ARFbs definition, the MP activator and the ARF2 repressor appear to have different preferred DNA binding sites. They share a novel inverted repeat (IR7-8) element but also have specific binding sites with different spacing and orientation of ARFbs (O’Malley et al., 2016). Here we undertook an extensive reanalysis of DAP-seq data using position weight matrix (PWM) as the DNA binding specificity model (Wasserman and Sandelin, 2004). PWMs represent a simple but efficient tool that captures the base preference at each position of the motif. PWMs give a score to any DNA sequence with zero for the optimal sequence and more negative scores as the sequence diverges from the optimum. The PWM score is then a quantitative value directly related to the affinity of the DNA molecule for the protein (Berg and von Hippel, 1987). Using PWMs, we establish differences between ARF2 and MP and show that they reliably identify a binding site syntax explaining their specificity. We further illustrate the predictivity of PWM as compared to consensus using binding assays and identify ARFbs configurations enriched in promoters of genes regulated by auxin.

## RESULTS

### ARF2 and MP have similar DNA binding sites but bind different genome regions

Using the published DAP-seq data (O’Malley et al., 2016), we first compared the sets of genomic regions bound by ARF2 and MP. Two regions were considered bound by both factors when they overlapped by at least 50% (see Methods). As expected for two TFs from the same family, there is a significant overlap and many regions are bound by both factors (Figure 1A). However, the large number of regions specifically bound by only one of them indicates a clear difference between ARF2 and MP DNA binding preferences (Figure 1A). This remains true even when focusing on regions bound with the highest confidence (top 10%, see Methods) by each of the factors (Supplemental Figure 1). We intended to explain these differences by characterizing ARF2 and MP DNA binding specificity. The examination of the DNA motif logo derived from regions recognized by ARF2 or MP monomers revealed only minor differences (Figure 1C). For both logos, the G[4] position corresponding to a direct protein-base contact in the ARF1 structure (Boer et al., 2014) is highly invariant. At positions [7,8] where the original ARFbs harboured TC (Guilfoyle et al., 1998), the preferred sequence is GG as recently proposed from PBM experiments (Boer et al., 2014; Franco-Zorrilla et al., 2014) but this preference is not as pronounced as in PBM-derived logos and sequence variations at these positions is tolerated.

**Figure 1:**
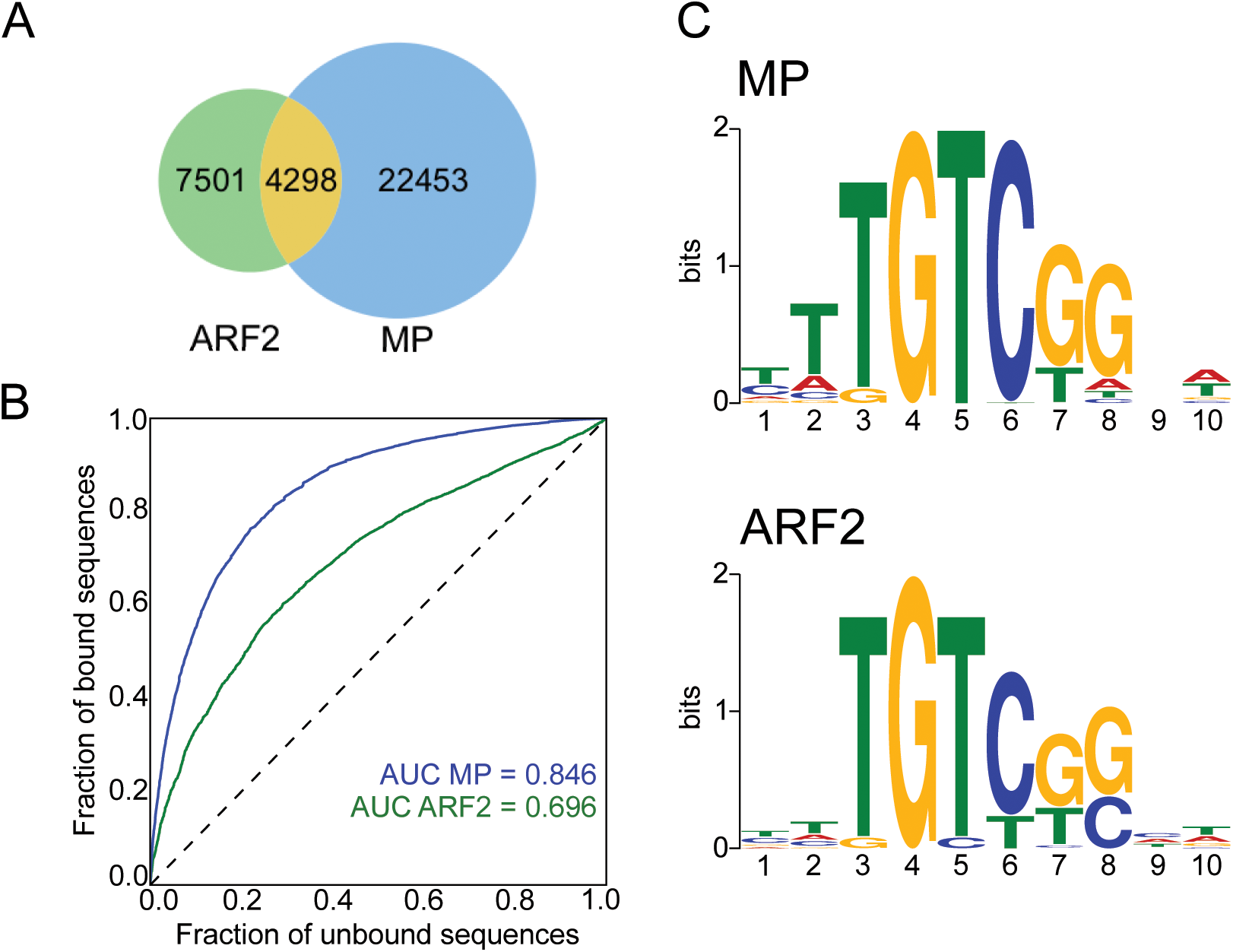
(**A**) Venn diagram of regions bound by ARF2 or MP in DAP-seq. (**B**) ROC curves and AUC values for MP and ARF2 PWM models. (**C**) Logo for MP and ARF2 PWM.

We built ARF2 and MP PWMs to model their DNA binding. We evaluated the prediction power of each PWMs using Receiver Operating Characteristics Area Under the Curve analysis (ROC-AUC or AUROC) (Hanley and McNeil, 1982) based on the ARFbs of best score present in each bound region (Figure 1B). Such analysis yields an AUROC value of 1 for a perfect model and 0.5 for a model with no predictive value. This analysis requires the generation of a negative set of regions for comparison. For this, we improved a previously designed tool, a negative set builder (Sayou et al., 2016), to extract from the Arabidopsis genome a set of non-bound regions with similar features as bound ones (size, GC content, genomic origin – see Methods). Based either on the full set of bound regions (Figure 1B) or only the 10% top ranked regions (Supplemental Figure 1), we found that MP model is highly predictive (AUROC=0.84) while ARF2’s has a lower performance (AUROC=0.69).

PWM models assume an additive contribution of each nucleotide position, a hypothesis that is not always true (Bulyk, 2002; Moyroud et al., 2011; Zhao et al., 2012; Mathelier and Wasserman, 2013). We used Enologos (Workman et al., 2005) to test for the presence of dependencies between some of the positions, particularly for positions [7,8] (Figure 1C) where mostly GG and TC doublets have been proposed so far. Enologos did not detect any dependency (Supplemental Figure 1) indicating that standard PWM can be adequately used. We also wondered whether the small differences between ARF2 and MP PWMs (as visible on their logos from Figure 1C) could contribute to their binding specificity. We thus tested the MP PWM on ARF2 regions and, conversely, ARF2 PWM on MP regions. The performance is indeed slightly weaker showing there is some specificity in the monomer PWM (Supplemental Figure 1). However, the very small difference suggests there must be other parameters explaining ARF2 and MP different specificities.

### ARF2 and MP prefer different binding site configurations

Published analyses (O’Malley et al., 2016) suggested that MP and ARF2 might differ in their preferred ARFbs dimeric configurations (ER, DR or IR, Figure 2A). We thus analysed the distribution of spacings between ARFbs using PWM models. To do this, a score threshold needs to be chosen above which transcription factor binding site (TFBS) are considered. As this threshold cannot be experimentally determined, we performed the analysis within a range of scores (from -8 to -13, -8 being of better affinity than -13). We studied the overrepresentations of all dimer configurations (DR, ER and IR) as compared to a negative set of regions. Overall, DR, ER and IR are more frequent in the ARF bound regions than in the negative set (Figure 2B, left panel), consistent with the higher density of ARFbs in these regions. We next estimated the overrepresentation of each particular configuration (ERn, DRn or IRn with the spacing n varying between 0 and 30 bp) within the whole population of configurations and normalized it to the equivalent parameter in the negative set of regions (Figure 2). For example, if, for a given value of n, DRn represents 10% of all configurations (ER/DR/IR with 0≤n≤30) in the positive set and only 2% in the negative one, DRn enrichment will be 5-fold.

**Figure 2:**
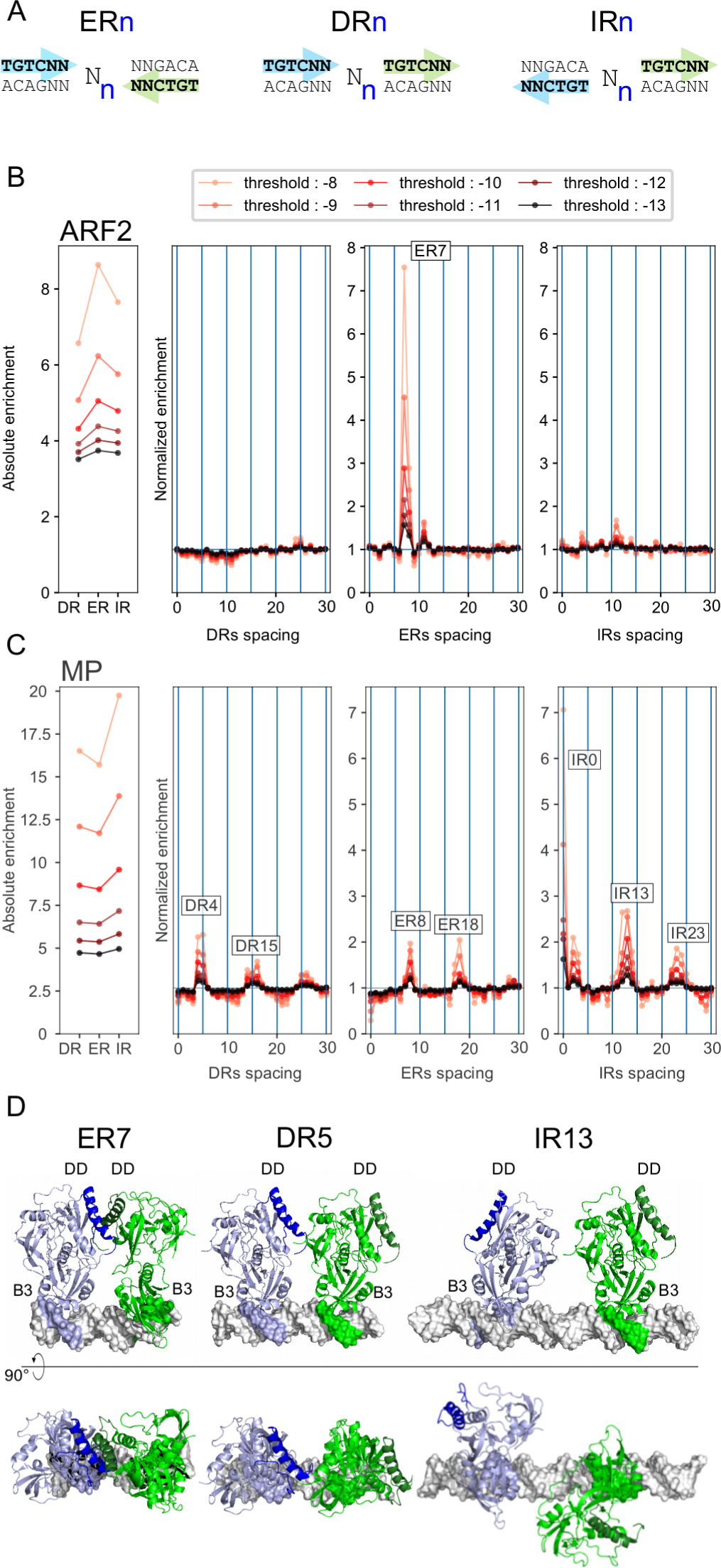
ARFbs configurations enrichment. (A) Definition of ERn, DRn and IRn. (B-C) Over-representation of dimeric ARFbs configurations in DAP-seq regions compared to an unbound set of sequences generated for ARF2 (B) and MP (C). The left panels quantify the absolute enrichment for all ERn, DRn and IRn (0≤n≤30) as compared to the background set. Right panels present the normalized enrichment for each ERn, DRn and IRn (see Methods). **(D)** Structural modelling of DR5 and IR13 ARF complexes based on ER7 ARF1 structure (PDB entry 4LDX) (Boer et al., 2014). Note the dimerization interface present on ER7 is absent in the two other configurations.

This analysis revealed a striking difference between ARF2 and MP. For ARF2, ER7-8 are the only overrepresented configurations whereas MP showed a wider range of preferred distances and configurations including DR4-5, DR14-15-16, DR25-26, ER7-8, ER17-18, IR0, IR3, IR12-13, IR23-24 (Figure 2B). Our results contrast with O’Malley’s where IR8 was the most overrepresented configuration for both factors (O’Malley et al., 2016). Since their result was obtained using a TGTC consensus as ARFbs definition, we repeated our analysis with TGTC (Supplemental Figure 2A). We still validated our result suggesting O’Malley et al. likely confused ER and IR. The MP graph (Figure 2B) suggests a periodicity of overrepresented distances every 10 bp, a hypothesis we confirmed by extending the distance window, revealing this trend for MP but not for ARF2 (Supplemental Figure 2B). Modelling of DR5 and IR13 protein/DNA complexes structures based on ARF1 crystallographic data (PDB entry 4LDX) clearly illustrates that these configurations are incompatible with the dimerization mode described for ER7 and could involve a different dimerization interface (Figure 2D).

### ARF2 and MP have different DNA binding syntax

We re-examined the Venn diagram from Figure 1A in the light of the identified preferred configurations. We separated ARF2 and MP bound regions in three sets: ARF2 specific, MP specific, ARF2/MP common regions. Because the two PWMs are very similar, we used the ARF2 matrix and performed the same analysis as in Figure 2 but on the three sets of regions (Supplemental Figure 3). DR4/5/15 and IR0/13 are overrepresented only in MP specific regions, ER7 in ARF2 specific regions and ER7/8 mostly in the MP/ARF2 common regions. Remarkably, MP-specific regions are even depleted in ER7/8 compared to the negative set of sequences because these elements are bound by both ARF2 and MP (Supplemental Figure 3). Plotting the frequency of a few selected configurations illustrates the group specific characteristics (Figure 3). We also used RSAT (Medina-Rivera et al., 2015) to search for other sequence features that could distinguish the three groups of regions. For ARF2-bound regions only, we found an enrichment for nine long AT-rich motifs similar to the one shown in Figure 3B. These motifs are found all along the bound regions (not shown). One example of enrichment of such a motif is illustrated in Figure 3B.

**Figure 3:**
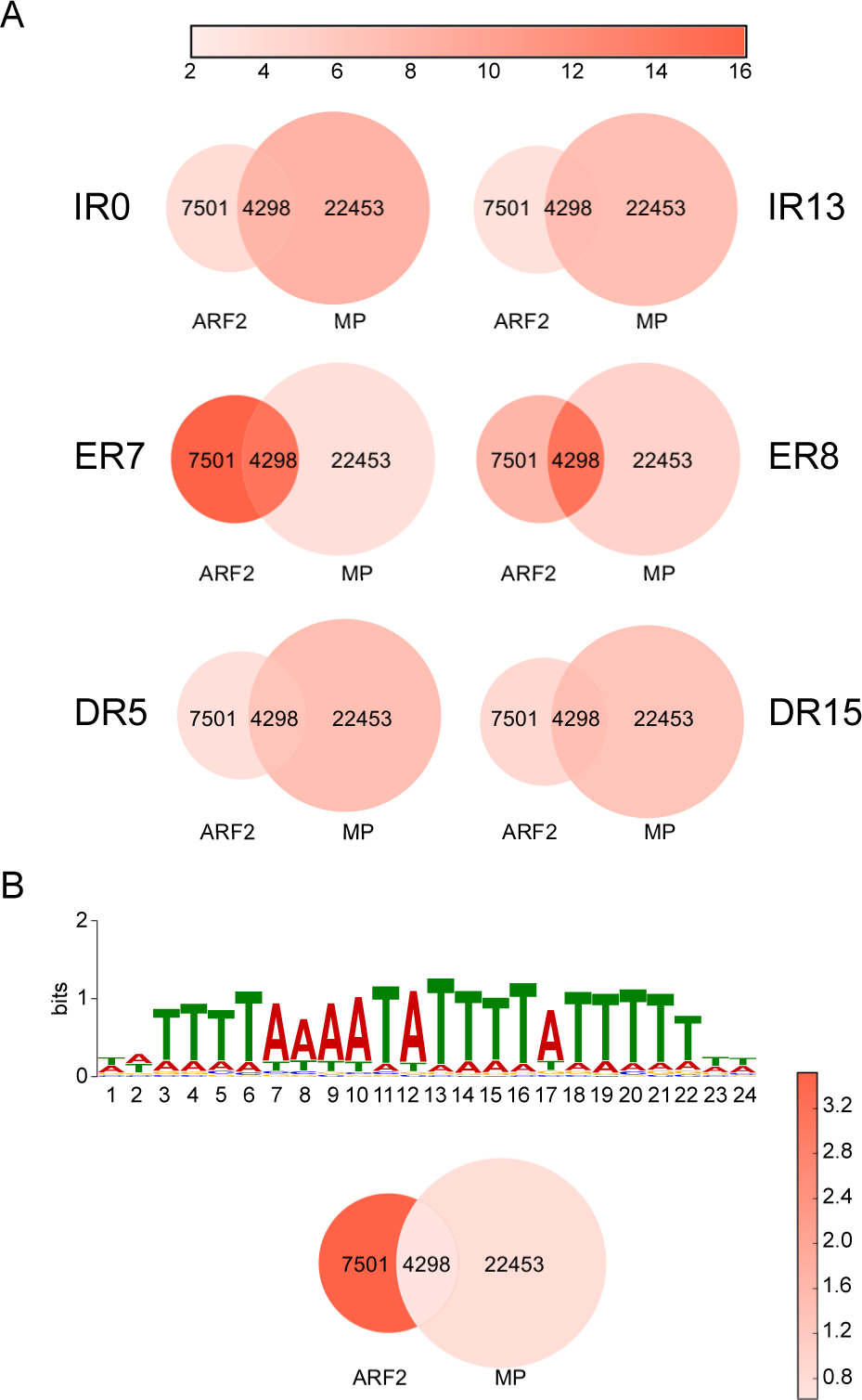
**(A)** Venn diagrams coloured according to the frequency (in %) of a few ARFbs conformations in MP-specific, ARF2-specific and MP/ARF2 common regions. (**B**) Fraction of regions containing at least one AT rich motif in MP-specific, ARF2-specific and MP/ARF2 common regions.

### Comparison between improved PWM models and consensus

We incorporated the ARF2 and MP specific features in new PWM-based models and tested their prediction power using AUROC. The improvement is marginal for MP but better for ARF2 (Figure 4C, AUROC for monomeric ARF2bs=0.69, for ER7/ER8 model=0.74). To illustrate the fundamental differences between PWM and consensus, we plotted the specificity (false positive rate) and sensitivity (true positive rate) parameters on the PWM ROC curve (Figure 4). For the monomeric ARFbs models, the TGTC consensus is poorly specific with almost 70% false positive rate. Conversely, TGTCGG or TGTCTC perform correctly but leave no freedom in terms of sensitivity and specificity: only the quantitative model allows to choose these parameters by adjusting the score threshold. For ARF2 ER7/8 dimeric models, using any of the three consensus is extremely stringent and detects very few sites in the positive set (at best 2.5% for TGTC) whereas the PWM model is again more versatile as it allows reaching the desired specificity/sensitivity combination by adjusting the score threshold.

**Figure 4:**
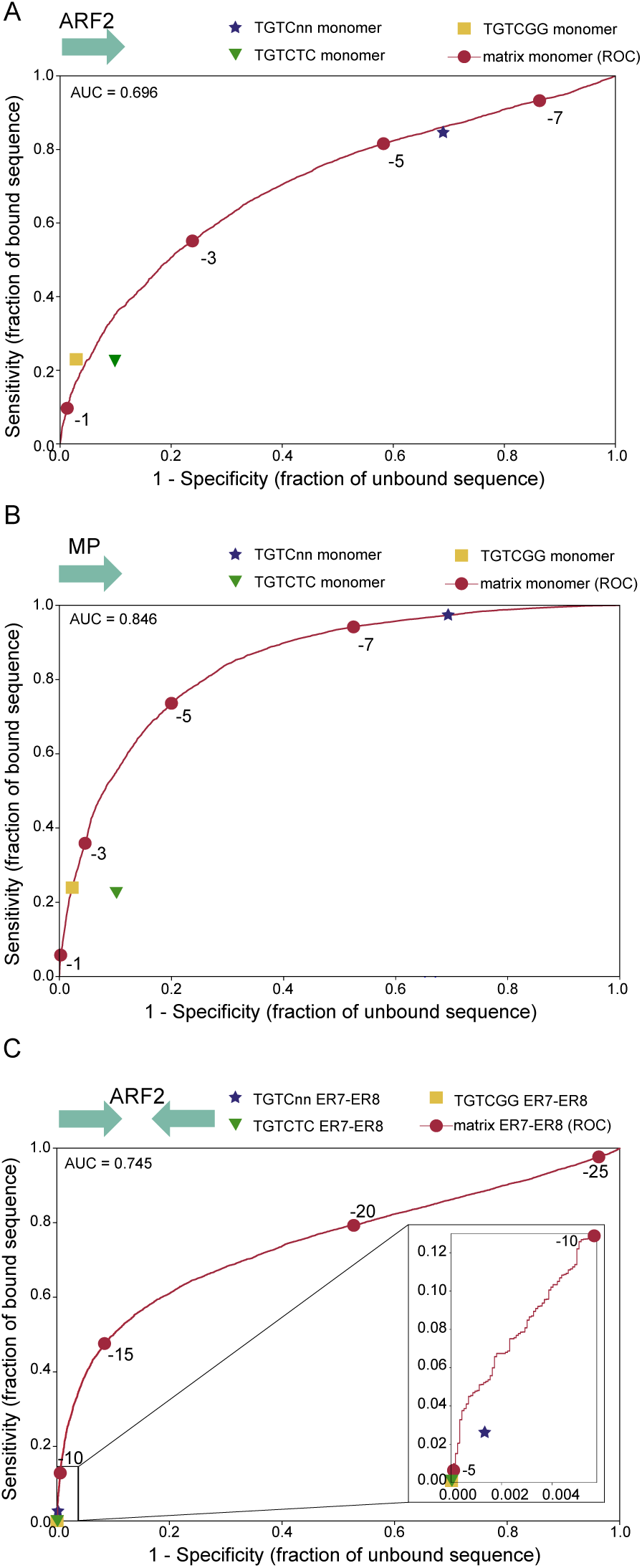
Comparison between PWM and consensus sensitivity and specificity. For all graphs, red dots correspond to score thresholds used to plot the PWM ROC curves. For consensus search, a sequence is considered positive for TGTC, TGTCGG or TGTCTC if this sequence is present at least once in the DNA region. The ER7-8 models were built as described in methods (**A**) ARF2 PWM and consensus on ARF2 bound regions. (**B**) MP PWM and consensus on MP bound regions. (**C**) ER7-8 PWM and consensus models on ARF2 bound regions.

DNA binding models are extremely useful to detect transcription factor binding site and challenge their role *in vitro* or *in vivo*. To scan individual sequences, PWMs are superior as they provide a quantitative information linked to TFBS affinity (Berg and von Hippel, 1987) and allow the detection of possible non-consensus sites of high affinity. We used our models to identify binding sites on the well-studied promoter of the *IAA19* gene ((Pierre-Jerome et al., 2016) and references therein). Scanning the *IAA19* promoter sequence with ARF2 and MP PWMs identified several ARFbs (Figure 5) including a high-scoring ER8 site bearing one non-canonical gGTCGG that lacks the TGTC consensus (Figure 5A). This site is located at the centre of a DAP-seq peak for MP and ARF2. We tested ARF binding to this particular ER8 element and tested the impact of the consensus presence on binding to ARF. For this, we restored the TGTC consensus for this non-canonical ER8 element and also created an artificial ER8 that has both TGTC consensus but suboptimal bases in other positions according to the PWM (Figure 5B). Strikingly, the optimised PWM score better predicts the binding than the presence of the consensus sequence: we observed intense binding on the non-canonical ER8, only a slight improvement when the consensus is restored and no binding on a consensus-bearing ER8 of low score (Figure 5C).

**Figure 5:**
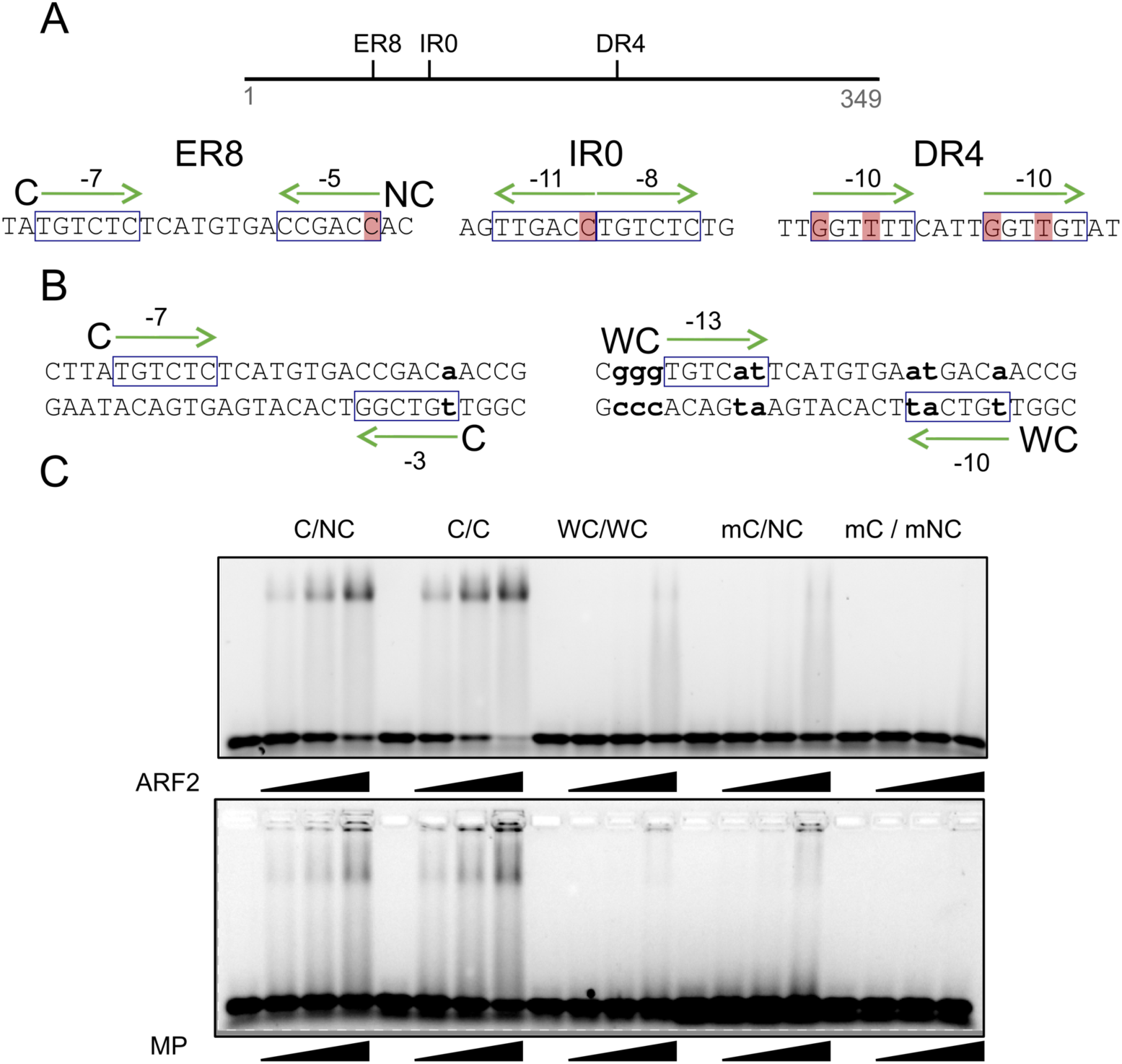
(A) Arabidopsis *IAA19* promoter with position, sequence and scores of ARFbs. (B) ER8 and its variants used in EMSA. (**C**) EMSA using ARF2 and MP proteins on probes described in B and two mutant probe controls with one (mC) or two mC/mNC sides of the ER8 mutated. ARF2 and MP are used at increasing concentrations: 0, 125, 250 and 500 nM. Colour shading indicates difference from consensus.

### PWM models reveal preferred ARFbs configurations in auxin regulated genes

We next tested the PWM models on *in vivo* data. ChIP-seq data are available for ARF6 and ARF3 (Oh et al., 2014; Simonini et al., 2017). However, no obvious ARFbs could be identified in any of these datasets. Testing ARFbs monomeric or dimeric models yielded a very poor AUROC value (0.61 for ARF6 and 0.58 for ARF3) suggesting that these data might not be adequate to evaluate our model. We also used the auxin responsive genes datasets derived from a meta-analysis of 22 microarray data (see Methods). We defined 4 groups of regions of either auxin induced or repressed genes with high or very high confidence (very-high confidence: 153 up regulated genes, 36 down regulated; high confidence: 741 up regulated, 515 down regulated, Supplemental File). We first analysed the 1500 bp promoters of the regulated genes compared to unregulated ones. This analysis revealed a mild but detectable over-representation of ER8 in up-regulated promoters (Supplemental Figure 4) as compared to unregulated ones and nothing in down-regulated genes.

Next, we tested whether more information could be extracted from these promoters if only the DNA segments bound by ARF in DAP-seq were analysed. We focused on auxin-induced genes and regions bound by the MP activator ARF because the mechanism of gene induction by auxin is well understood, while repression by auxin and the role of repressor ARFs such as ARF2 is less clear. We therefore compared MP-bound regions present in regulated versus non-regulated promoters. We observed that the over-representation of ER8 and IR13 is higher in auxin upregulated genes than in non-regulated ones (Figure 6A-B). This is particularly striking for the high-confidence auxin induced genes even if this list likely also contains indirect ARF targets (Figure 6A). We tested MP binding to the IR13 probe and observed a strong and well-defined MP/IR13 complex (Figure 6C), similar to those obtained with ER7/8 probes. The IR0 element, also enriched in MP-bound DAP-seq regions but not in auxin-regulated promoters, gives a weaker smeary band. For auxin repressed genes, two configurations (ER18 and IR3) are more overrepresented in the MP-bound regions from promoters of downregulated genes than for non-regulated genes (Supplemental Figure 5). This might indicate that some ARFbs configurations could be involved in repression by auxin but this attractive hypothesis clearly requires to be tested with additional experiments.

**Figure 6:**
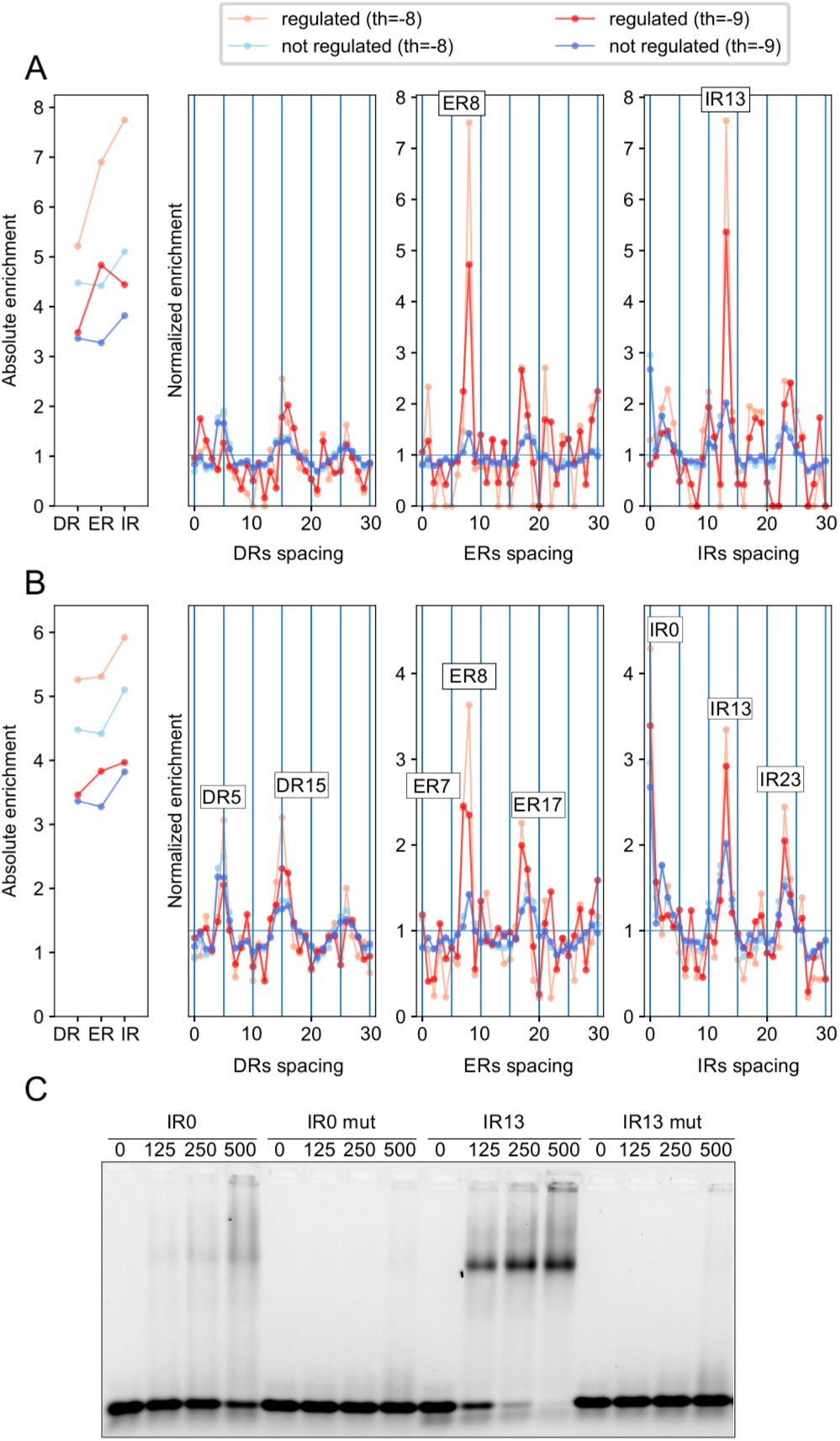
Spacing enrichment in promoter regions bound by MP were analysed in auxin up-regulated very high-confidence (**A**) or high-confidence genes (B) (red colours) and non-auxin-regulated genes (**B**) (blue colours) at two different score thresholds. The enrichment of ER7/8 and IR13 is increased for genes of the very high confidence auxin upregulated gene list. (C) EMSA showing the binding of MP to IR0 and IR13 motifs and the corresponding control mutant probes. MP is used at increasing concentrations: 0, 125, 250 and 500 nM.

## DISCUSSION

### PWM versus consensus for auxin responsive elements

A key question in auxin biology is how the structurally simple molecule evokes such diverse responses. Transcription Factors of the ARF family are the main contributors that diverge auxin response. Predictive tools to infer the presence of ARF binding sites in regulatory regions are essential both for functional and evolutionary analyses. Most studies so far have used TGTC-containing consensus sequences as a tool to detect ARFbs (Berendzen et al., 2012; O’Malley et al., 2016; Zemlyanskaya et al., 2016). Here we built PWM-based models and showed that they provide a greater versatility than consensus sequences as they allow adjusting sensitivity and specificity. Even if a TGTC consensus is perfectly suitable to detect the over-representation of some configurations (such as ER7-8 for ARF2 and MP)(O’Malley et al., 2016) (Supplemental Figure 2), it cannot be used to search regulatory regions because of its lack of specificity when used as monomer and its extremely low sensitivity when used as ER7/8 dimer (Figure 4). We illustrated on a chosen example (the *IAA19* promoter) that a site can be bound without a TGTC consensus and not necessarily bound even when the consensus is present (Figure 5). The non-canonical ER8 site we detected was challenged and functionally validated by studies in yeast (Pierre-Jerome et al., 2016).

Even if more elaborate models exist (Mathelier and Wasserman, 2013), PWM have emerged as the simplest and most performant models. Still, we were surprised that, in a DAP-seq context where no other parameters (such as cofactors, histones or chromatin accessibility) should influence TF/DNA binding, the PWM models could not reach better AUROC values especially for ARF2. We have tried models that integrate the DNA shape feature (Mathelier and Wasserman, 2013) but they did not significantly improve the prediction power (data not shown). The newly identified sequences with stretches of As and Ts (Figure 3B) were not easy to integrate in improved models but might affect the overall context of ARF2 binding sites and contribute to ARF2 specific regions. This finding is reminiscent of the family of AT-rich motifs found as overrepresented in promoters of auxin responsive genes (Cherenkov et al., 2018). These elements were mostly found in ARF2-binding regions and they were more associated with down-regulation than with up-regulation. More studies are needed to elucidate their role.

### ARF2 versus MP

ARFs exist as activators and repressors (Dinesh et al., 2016). Affinity measurements on a few DNA sequences *in vitro* (the molecular caliper model) and consensus-based search in genome-wide binding data both indicate that activator ARF MP and repressor ARF (ARF1 and ARF2) might have different preferences for ARFbs configurations (Boer et al., 2014; O’Malley et al., 2016). But one study examined only a few ER elements (Boer et al., 2014) whereas the other did not recover the long known ER7/8 elements and instead proposed IR7/8 (O’Malley et al., 2016). Using PWM-based models and re-analysing DAP-seq data, we confirmed that MP and ARF2 have a similar monomeric binding site but differ in the syntax of binding sites (combinations of binding sites of ARF monomers) they recognize: ARF2 prefers ER7/8 while MP has a much wider range of preferences. For ER motifs (face to face DBDs), our results extend the molecular caliper model (Boer et al., 2014) at the genome-wide level with some larger spacings. Moreover, MP has wider syntax than ARF2 as it also includes enriched DR and IR motifs. Such findings cannot be accommodated with the molecular caliper model as they involve different orientations of the two DBDs than in ER (head to tail for DR and tail to tail for IR). As previously reported (O’Malley et al., 2016), MP shows an increased binding frequency every 10 bp for all DR, ER and IR enriched configurations. Because this spacing corresponds to a DNA helix turn, we can imagine that this configuration allows interaction between ARF proteins on the same side of the DNA. 3D modelling using the published ARF1 structure indicates that these interactions are unlikely to involve the same dimerization surface as for ARF1 (Figure 2D). The proximity of some ARF DD domains in 3D, combined with possible flexibility of ARF DBD suggest that these proteins might have evolved different dimerization modes with the same protein domain. Confirming this hypothesis will await their structural characterisation. An alternative hypothesis is that the PB1 oligomerization domain contributes to stabilize the MP binding to preferred motifs but this also remains to be tested. However, it should be also noted that a preference for 10-bp spaced binding sites does not necessarily implies the presence of protein-protein contacts. Indeed, it has been shown that the binding of a first protein in the DNA major groove favours the binding of a second one at a 10 bp distance through allosteric changes in DNA conformation (Kim et al., 2013). This mechanism could also be at work for ARF DNA binding.

It is interesting to note that ER7-8 is bound by both ARF2 and MP whereas some configurations such as DR5 or IR13 are more specific to MP. If repressor ARFs act by competing with activator ARFs for ARFbs (as proposed in Vernoux et al., 2011), this competition will therefore depend on the nature of ARFbs (shared between activators and repressors or specific to only one class of ARFs). Some sites such as DR5 might be less subjected to competition therefore influencing their activity as reporter for auxin-dependent transcriptional activity (Ulmasov et al., 1997; Liao et al., 2015). Extending this type of analysis to all members of the ARF family should indicate whether ARF from a given class (A, B or C) (Finet et al., 2013) have a stereotypic behaviour or whether there is also a diversity of properties within the class A ARF, for example. Such differences would help explaining how a single auxin signal can trigger different responses depending on the cell type where it is perceived (provided different cell types express different sets of ARF proteins). *In vivo*, other parameters will also play an important role for the response to auxin such as the ARF interaction partners (Mironova et al., 2017) and chromatin accessibility.

### ARF binding versus auxin regulation

The analysis of auxin-induced genes using PWM models identified only a small over-representation of ER8 (Supplemental Figure 4), a motif shared by ARF2 and MP. As we anticipated that ARFbs might be diluted in whole promoter sequences, we collected the set of DNA regions present in promoters from auxin-induced genes that are also bound by MP in DAP-seq and compared it to MP-bound promoter regions from non-auxin-regulated genes. This analysis confirmed the overrepresentation of ER8 in auxin-induced genes but also identified IR13 as enriched motifs (Figure 6). IR13 is a novel element, well bound by MP *in vitro* that now requires *in planta* characterization. It is not enriched in ARF2-bound regions suggesting it will likely be insensitive to competition by repressor ARF2. We also characterized auxin repressed gene. Whether repression directly involves ARFs is not known. Promoter analysis did not reveal any motif enrichment but the intersection with MP-bound regions showed ER18 and IR3 over-representation (Supplemental Figure 5). Again, functional analysis of such motifs *in planta* will be important in the future. We anticipate that the strategy we designed here (combining DAP-seq data with expression studies) is a very general method to increase the signal/noise ratio in regulatory regions and better detect binding sites involved in regulation. DAP-seq is a powerful technique but it suffers from giving access to DNA that might never be accessible in the cell. The combination with differential expression studies (+/- a stimulation or +/- a TF activity) will be a powerful way to narrow down the number of regions examined and extract functional regulatory information.

## METHODS

### Bio-informatic analyses

The TAIR10 version of Arabidopsis genome was used throughout the analyses. The DAP-seq peaks were downloaded from http://neomorph.salk.edu/PlantCistromeDB. We sorted the peaks (200 bp) extracted from narrowPeaks file accordingly to their Q-value. An ARF2-bound region was considered to overlap with an MP-bound region if the overlap exceeded 100 bp. We used the Bedtools suite to assess the overlaps and retrieve genome sequences. The PWM were generated using MEME Suite 4.12.0 (Bailey et al., 2009) on the 600 top peaks of ARF2 and MP according to the Q-value.

ROC-AUC analysis: performing a ROC analysis requires a background set of unbound genomic regions. This set was built with a Python script that takes a *bed* file of bound genomic regions as input and randomly selects in the Arabidopsis genome regions of same size, similar GC content and with similar origin (intron, exon or intergenic).

To search for dependencies between positions of the ARF PWM, we used the sequence alignment inferred by MEME Suite (Bailey et al., 2009) to build a PWM and used it as input for Enologos selecting the option “mutual info” (Workman et al., 2005).

#### Analysis of ARFbs configurations

The absolute enrichment (A) for each type of configuration (DR, ER, IR) was calculated as the ratio between the total number of sites in each configuration C in the bound set of regions divided by the same number in the background set. Such calculations were done for different score thresholds and normalized by the ratio between the total number of monomeric sites (BS, with no threshold applied) in the foreground and in the background to account for the different sequence sizes of the two sets. *S_max_*stands for the maximum spacing.

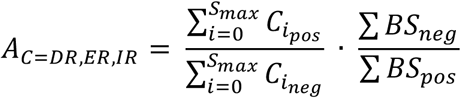

For the normalized enrichment, we inventoried all the dimer configurations made of two monomeric ARFbs with scores above the chosen threshold. We then calculated the frequency (f) of each particular conformation (DRn, ERn or IRn with 0≤*n*≤*S_max_*) among all dimeric sites in the positive set of bound regions and in the background set.

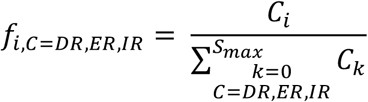

The normalized enrichment (*N*) shown in figure 2 corresponds to the ratio between frequencies in the positive set and in the negative set for a given spacing.

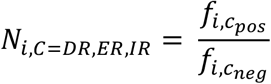

To illustrate the enrichment of a few chosen motifs (DR4-15, ER7-8, IR0-13), we identified all sequences displaying a potential binding site with a score higher than a - 8 threshold. Next, we plotted the % of regions displaying a given motif in the Venn diagram regions. The same was done for AT-rich motifs with a score threshold for each AT-rich PWM of a score -10.

The ER7/ER8 PWM for ARF2 was built from the ARF2 monomer PWM. Both ARF2 bound and unbound sets of regions were scanned with these two PWM and the best score given to each region by either ER7 or ER8 was used to plot the ROC curve. For the analysis of specificity and sensitivity of TGTC-containing consensus sequences, we analysed each region for the presence or absence of ER7 or ER8 consensus (TGTCNN-7/8N-NNGACA, TGTCGG-7/8N-CCGACA, TGTCTC-7/8N-GAGACA). A region was scored positive when containing at least one site.

For the analysis of auxin regulated promoters, we used 1500 bp upstream of the first exon of each gene. All DAP-seq regions overlapping with the promoters were then selected for analyses.

The major scripts used are available on github: https://github.com/Bioinfo-LPCV-RDF. The frequency matrices used to infer PWM can be downloaded on https://github.com/Bioinfo-LPCV-RDF/Scores.

#### Selection of auxin regulated genes

We selected auxin regulated genes over twenty-two publicly available gene expression profiling datasets from the GEO database (Supplemental File 1). The datasets were generated on seedlings or roots of *A. thaliana* with different auxin concentrations and times of exposure to auxin (explored in Zemlyanskaya et al., 2016). Differentially expressed genes were defined as those expressed at least 1.5 times higher (lower) after auxin treatment comparing to control, with FDR adjusted p-value < 0.05 (Welch t-test with Benjamini-Hochberg correction). We compiled four groups of auxin-regulated genes: induced or repressed genes with high or very high confidence (Supplemental File 1). High confidence genes: 741 auxin up-regulated and 515 down-regulated genes significantly (more than 1.5-fold, FDR adjusted p < 0.05) changed their expression after auxin treatment in two or more datasets. Very high confidence genes: 153 auxin up-regulated and 36 down-regulated genes significantly changed their expression in four or more datasets.

### Expression and purification of recombinant proteins

ARF2 and ARF5 coding sequences were cloned into pHMGWA vectors (Addgene) containing N-terminal His-MBP-His tags. His-MBP-His-tagged ARF proteins were expressed in *E. coli* Rosetta2 strain. Bacteria cultures were grown in liquid LB medium to an O.D_600 nm_ of 0.6-0.9. Protein expression was induced with isopropyl-β-D-1-thyogalactopiranoside (IPTG) at a final concentration of 400 µM. Protein production was done overnight at 18°C. Bacteria cultures were centrifuged and the resulting pellets were resuspended in Lysis buffer (Tris-HCl 20 mM pH 8; NaCl 500 mM; Tris(2-carboxyethyl) phosphine (TCEP) 1 mM for ARF2 and Tris-HCl 20 mM pH 8; NaCl 500 mM; EDTA 0.5 mM; PMSF 0.5 mM; TCEP 1 mM; Triton 0.2% (w/v) for ARF5) with EDTA-free antiprotease (Roche) for sonication. Proteins were separated from the soluble fraction on Ni-sepharose columns (GE Healthcare) previously equilibrated with the corresponding Lysis buffer. Elutions were done with Imidazole 300 mM diluted in the corresponding Lysis buffer.

### Electrophoretic Mobility Shift Assays (EMSAs)

DNA-protein interactions were characterized by EMSAs. ER8 binding site was isolated from Arabidopsis *IAA19* promoter and ER8 variant sequences are given in Supplemental Table 1. IR0 and IR13 sequences were artificially designed with TGTCGG consensus sites (Supplemental Table 1). EMSA DNA probes were prepared from lyophilized oligonucleotides corresponding to the sense and antisense strands (Eurofins). Oligonucleotides for the sense strand presented an overhanging G in 5’ for DNA labelling. Sense and antisense oligonucleotides were annealed in Tris-HCl 50 mM; NaCl 150 mM. The annealing step was performed at 98°C for 5 minutes, followed by progressive cooling overnight. Annealed oligonucleotides, at a final concentration of 200 nM, were incubated at 37°C for 1 hour with Cy5-dCTP (0.4 μM) and Klenow enzyme in NEB2 buffer (New England Biolabs). The enzyme was inactivated by a 10-minutes incubation at 65°C. Oligonucleotides were conserved at 4°C in darkness. EMSAs were performed on agarose 2% (w/v) native gels prepared with Tris-Borate, EDTA (TBE) buffer 0.5 X. Gels were pre-run in TBE buffer 0.5 X at 90 V for 90 minutes at 4°C. Protein-DNA mixes nonspecific unlabelled DNA competitor (salmon and herring genomic DNA, Roche Applied Science; final concentration 0.045 mg/ml) and labelled DNA (final concentration 20 nM) in the interaction buffer 25 mM HEPES pH 7.4; 1 mM EDTA; 2 mM MgCl2; 100 mM KCl; 10% glycerol (v/v); 1 mM DTT; 0.5 mM PMSF; 0.1% (w/v) Triton. Mixes were incubated in darkness for 1 hour at 4°C and next loaded in the gels. Gels were run for 1 hour at 90 V at 4°C in TBE 0.5 X DNA-protein and bindings were visualized on the gels with Cy5-exposition filter (Biorad ChemiDoc MP Imaging System).

## AUTHOR CONTRIBUTION

FP and RD designed and supervised the project, AS, JL, AB and VM performed the bioinformatic analyses, RMA and TVP performed the biochemical experiments, all authors discussed the results, FP wrote the manuscript with the help of AS, RD, TV, RMA and VM.

## ACKNOWLEDGEMENTS

We thank Anthony Mathelier for discussions, Line Andresen and Chloe Zubieta for critical reading of the manuscript and David Mast and Laura Grégoire for implication in early stage of this work.

## SUPPLEMENTAL INFORMATION

Supplemental Information is available at Molecular Plant Online.

**Supplemental Figure 1:**
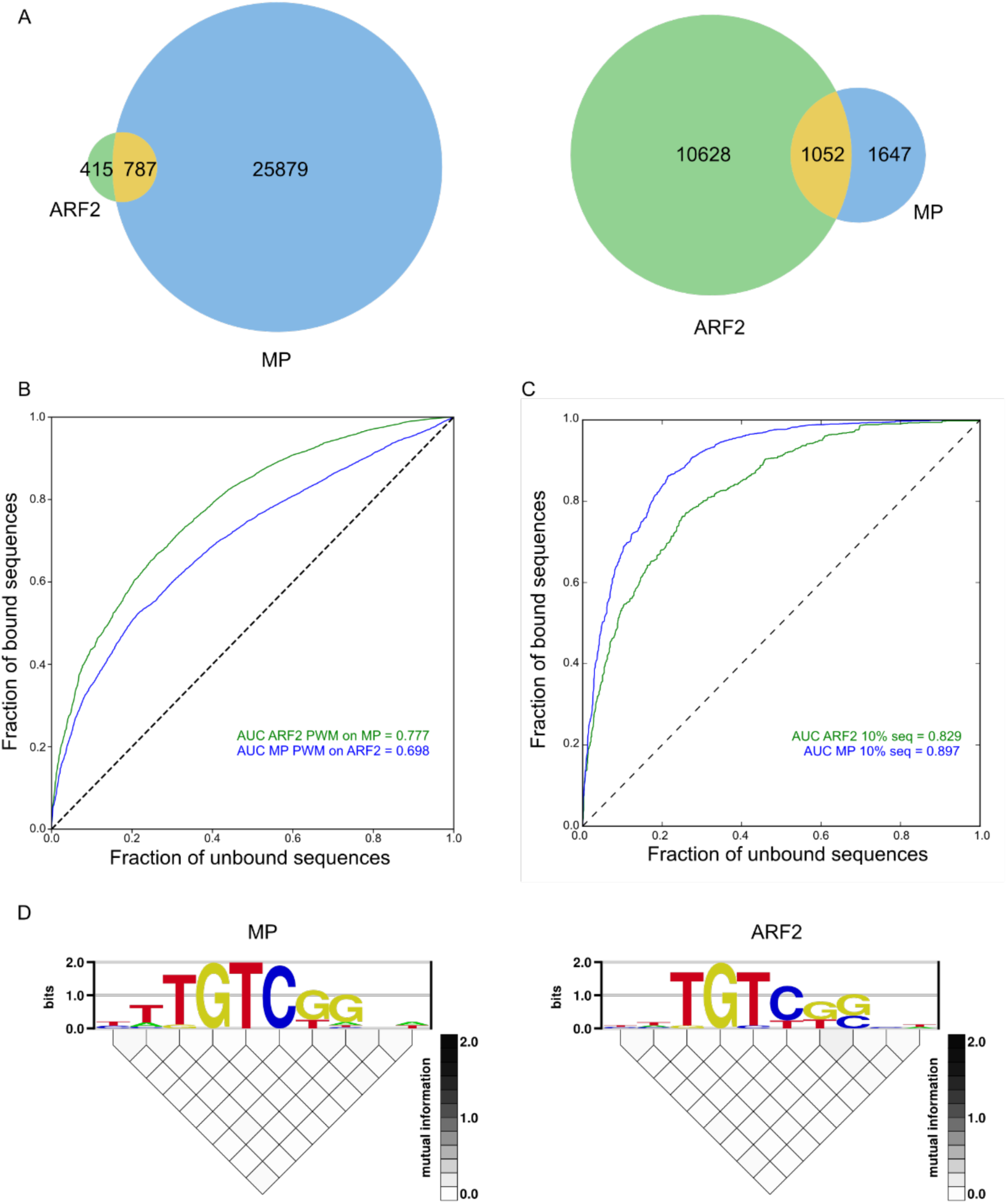
**(A)** 2 Venn diagrams with the 10% top bound regions for ARF2 against all MP regions and the 10% top bound regions for MP against all ARF2 regions. This shows that there are regions specifically bound by a single factor even in the highest confidence regions **(B-C)** ROC curves with ARF2 PWM on MP bound regions and MP PWM on ARF2 regions. AUROC value decrease slightly as compared to Figure 1 **(D)** Enologos analysis of MP and ARF2 motifs (1). No dependency between nucleotide position is detected.

**Supplemental Figure 2:**
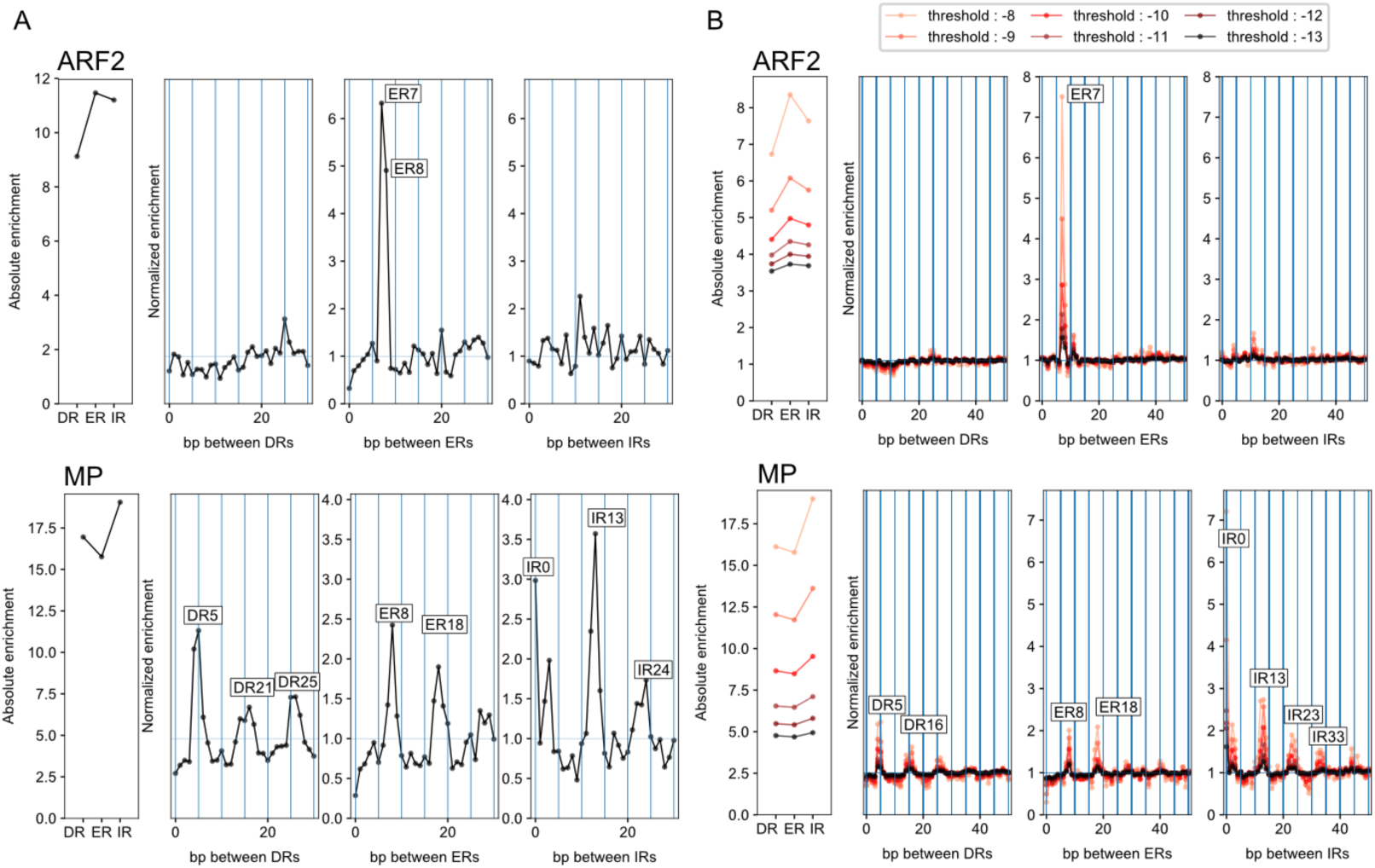
**(A)** Enrichment of spacings between TGTC **(B)** Spacing enrichment for DRn, ERn and IRn for 0≤n≤50. Threshold indicates the PWM score threshold value used for ARFbs detection

**Supplemental Figure 3:**
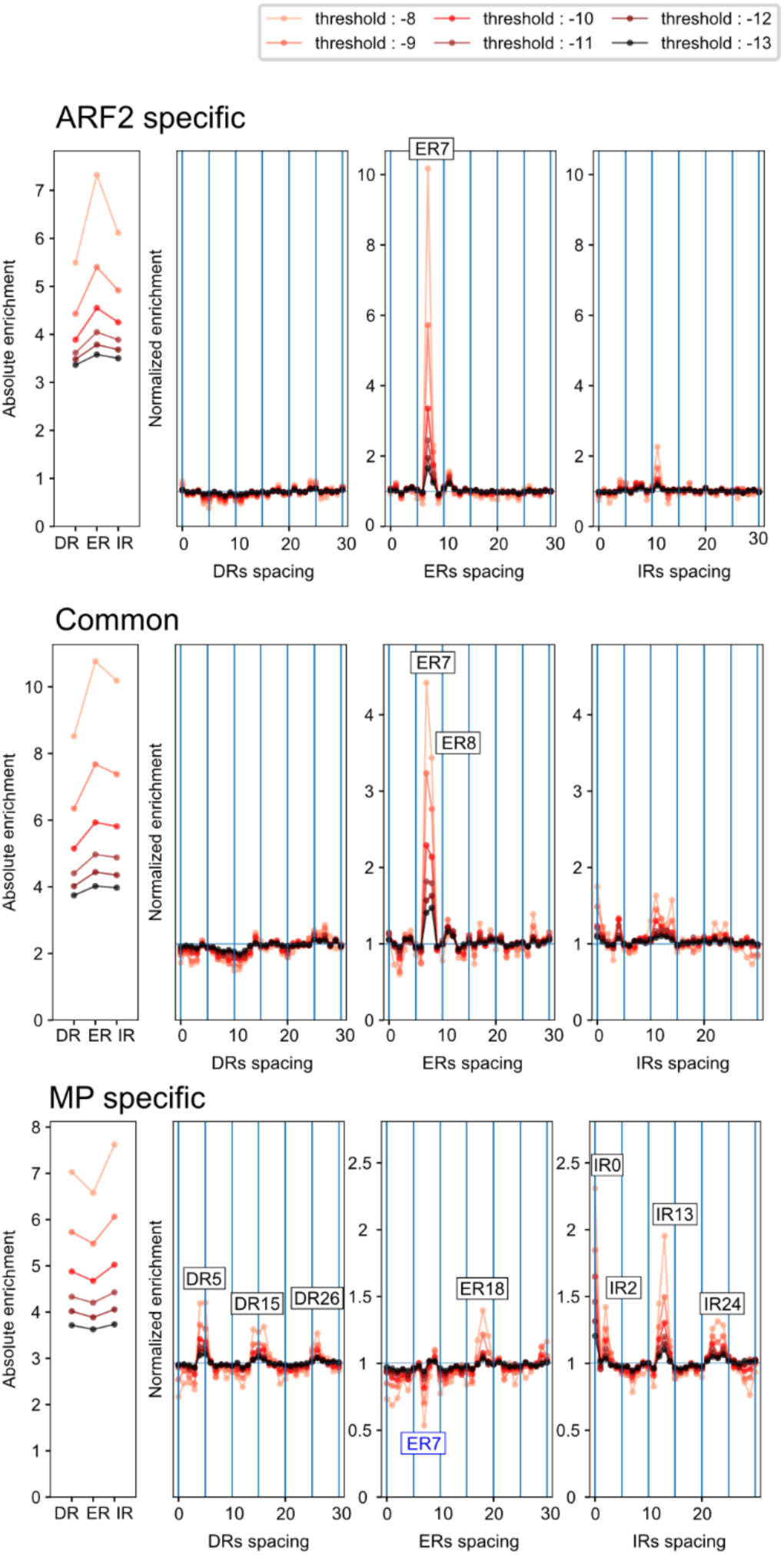
Spacing enrichment in MP-specific, ARF2-specific and MP/ARF2 common regions, compared to unbound sets of sequences. Threshold indicates the PWM score threshold value used for ARFbs detection. Note ER7 is depleted in MP-specific bound regions.

**Supplemental Figure 4:**
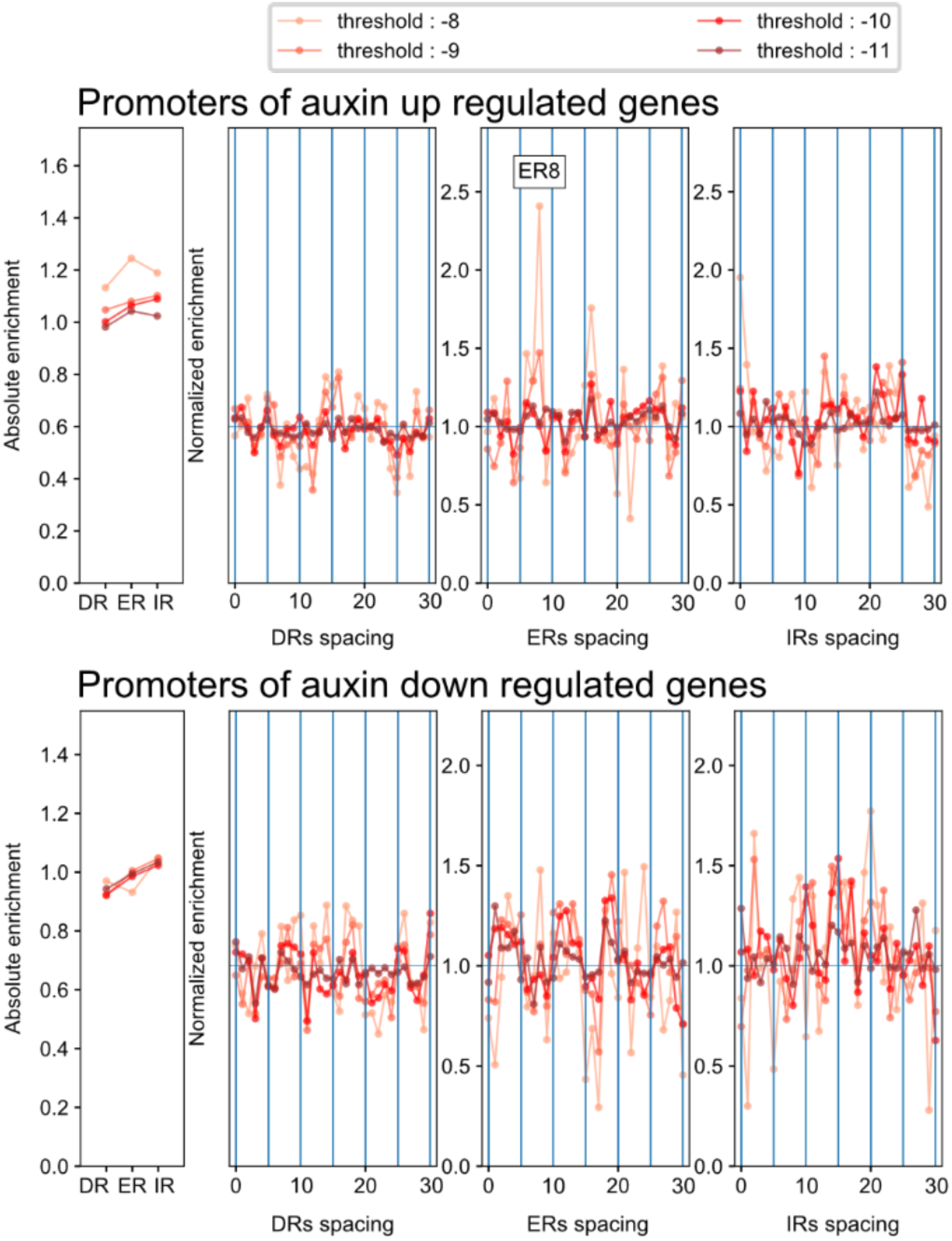
ARFbs over-represented conformations in the promoters of the auxin up-regulated genes (upper panel) or the down-regulated genes (lower panel) We used very high and high confidence genes and compared to auxin insensitive gene promoters. Threshold indicates the PWM score threshold value used for ARFbs detection

**Supplemental Figure 5:**
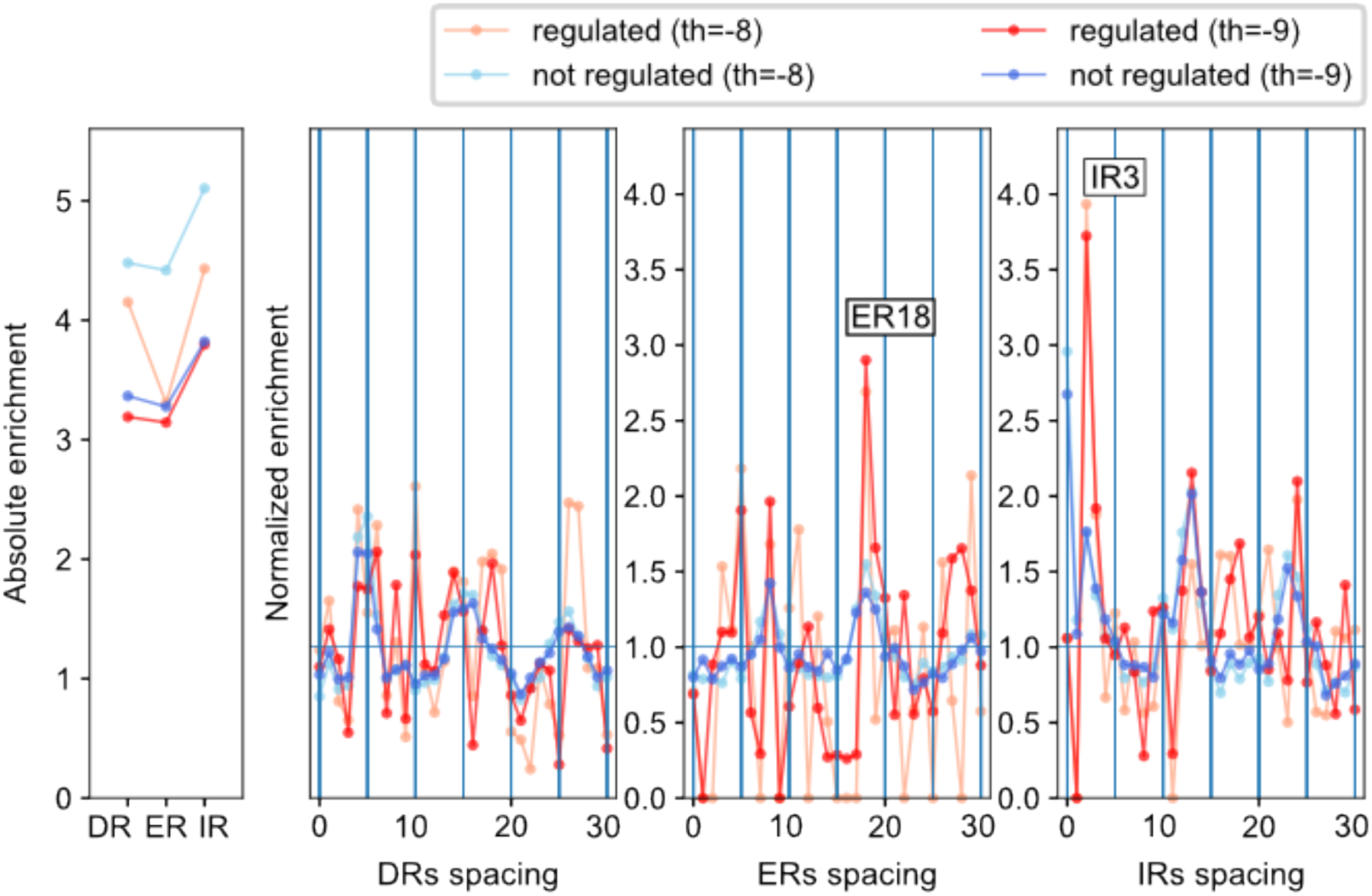
Promoter regions bound by MP were analysed in down-regulated (red colours) and non-regulated genes (blue colours) in high confidence gene lists (Supplemental file 1). The regions not bound by MP from auxin insensitive promoters were used as background. Threshold indicates the PWM score threshold value used for ARFbs detection

**Supplemental Table 1.**
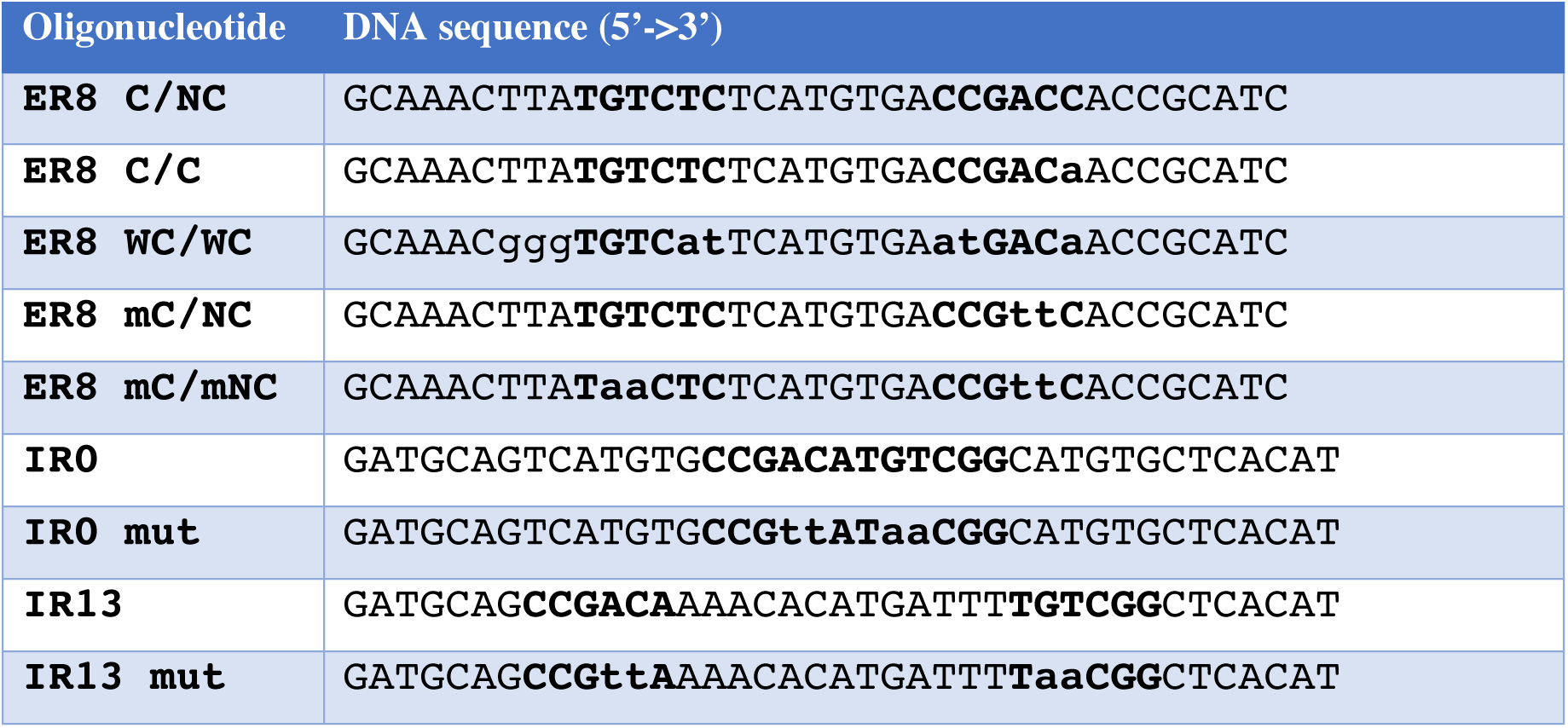
Sequences of DNA probes for EMSAs. Bold letters show ARF binding sequence. Lower case letters indicate the nucleotides variation introduced.

